# Scalable electron tomography for connectomics

**DOI:** 10.1101/2024.06.05.597487

**Authors:** Aaron T. Kuan, Sébastien Phan, Keun-Young Kim, Mason Mackey, Minsu Kim, Steven T. Peltier, Mark Ellisman, Wei-Chung Allen Lee

## Abstract

We demonstrate limited-tilt, serial section electron tomography (ET), which can non-destructively map brain circuits over large 3D volumes and reveal high-resolution, supramolecular details within subvolumes of interest. We show accelerated ET imaging of thick sections (>500 nm) with the capacity to resolve key features of neuronal circuits including chemical synapses, endocytic structures, and gap junctions. Furthermore, we systematically assessed how imaging parameters affect image quality and speed to enable connectomic-scale projects.

## Introduction

Connectomic datasets have helped us understand how neurons are connected and process information in the nervous system. For example, connectomics has recently provided insights into how neuronal circuit connectivity supports pattern association and memory^1,2^, decision-making^3,4^, and motor control^5,6^. Connectomic data have primarily been generated with electron microscopy (EM) due to its high spatial resolution and comprehensive labeling, and automated EM systems have been developed to increase imaging throughput^7–9^. For example, the “GridTape” approach uses automated, tape-based thin-sectioning and sample handling, which enables large-scale 3D TEM imaging^10^. However, large-scale, volume EM methods tend to image at modest resolutions (∼4-8 nm/pixel in x,y and ∼8-50 nm in z) to increase imaging speed, which resolves neuronal processes and chemical synapses, but may miss other key cellular components such as electrical synapses (gap junctions) and other mechanisms of cell-to-cell communication^11,12^.

Structural correlates of such signaling mechanisms can be resolved via electron tomography (ET), which combines images of the sample at multiple tilt angles (**Fig. 1a, b**) to achieve high resolutions within thicker sections (200-1000 nm). ET can resolve molecular details of biological structures in tissue samples^13–19^, and reveal atomic structures of single particles. However, ET is typically very slow due to the need to acquire many images of the same area, and the associated additional sample handling.

**Figure 1.**
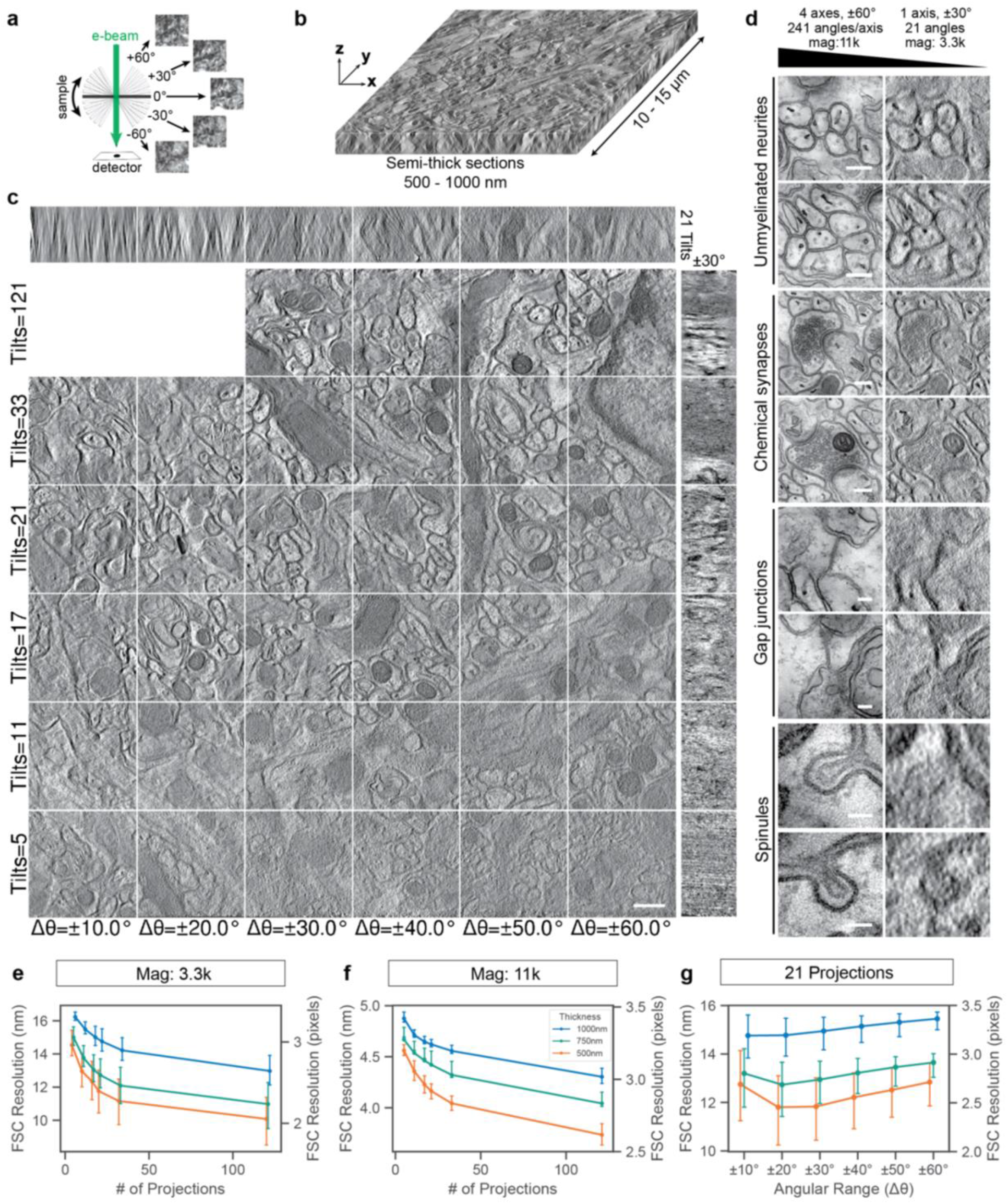
Overview of scalable electron tomography (ET). **(a)** Schematic of ET data acquisition. 2D image projections (images, right) of the sample at a range of tilt angles (black and gray bars) are acquired using a transmission electron microscope. **(b)** 3D volume of the sample (tomogram) computationally reconstructed from multiple 2D tilt projections. **(c)** A comparison of ET images of a 750 nm semi-thick section from the molecular layer of an adult mouse cerebellar cortex. Different subportions of the image matrix show data reconstructed using different tilt ranges (x-axis) and number of tilt angles (y-axis, evenly sampled over the tilt range). X-Z *(top bar)* and Y-Z *(right bar)* virtual slices through the tomographic volume are shown from different operating points. Note, differences in data quality with different tilt ranges and angles. Scale bar, 500 nm. **(d)** Biological features including unmyelinated neurites, chemical synapses, gap junctions, and spinules from tomographic volumes acquired with full (left column) and limited-tilt (right column) tomography. Scale bars: Unmyelinated neurites and Chemical synapses, 250 nm; Gap junctions, 100 nm; Spinules, 50 nm. **(e)** Plot of Fourier shell correlation (FSC) resolution as a function of the number of imaged projections equally spaced across an angular range (Δθ) of ± 30º and at a magnification (mag) of 3.3k. Left y-axis indicates FSC resolution in nm, whereas the right y-axis indicates resolution in pixels. Colored lines denote nominal section thickness (color legend in (f), blue: 1000 nm, green: 750 nm, orange: 500 nm). Error bars indicate mean ± IQR. **(f)** Similar to (e) at 11k mag. **(g)** Plot of FSC resolution as a function of angular range (Δθ) for a constant number of imaged projections (n = 21). Colored lines denote nominal section thickness as in (e,f). Note, angular range corresponding to the highest resolution is less than ±30º and depends on thickness. Error bars indicate mean ± IQR.

Here, we increased imaging throughput of ET by using angular subsampling to reduce imaging and sample handling time, while still retaining sufficient image quality to resolve biological structures of interest. We: (1) establish that intermediate-voltage ET can image semi-thick (500-1000 nm) sections with sufficient resolution to identify connectomic features, including gap junctions and spinule-like structures; (2) determine limited-tilt tomography parameters appropriate for high-throughput connectomic imaging; and (3) propose workflows for data acquisition, alignment and segmentation of serial section ET. These results suggest that subsampled, limited-tilt serial section ET is a robust, high-throughput, and sample-preserving approach for multiscale connectomic imaging.

## Results

We first sought to parameterize ET acquisition by known factors that limit throughput, including the number of rotational axes, tilts or projections, range of angles, and magnification. These are critical parameters for improving acquisition speed because physical movements and settling of the sample limit acquisition throughput. High-resolution, multiple-tilt ET can use multiple rotational axes, with ±60º tilt angle range (Δθ) distributed evenly over hundreds of tilts per axis. Here, we refer to using four rotational axes, Δθ = ±60º, with 241 tilts per axis as “fully-sampled” ET. Our goal was to identify a “limited-tilt” regime that still resolves neuronal processes and synapses for connectomics.

Using a 300 kV TEM (FEI Titan Halo) equipped with a 8k × 8k direct electron detection camera (DE-64), we acquired tomograms from 750 nm semi-thick sections from the molecular layer of mouse cerebellar cortex at magnifications of 3,300× or 11,000×. We performed tomographic reconstructions using different subsets of the projections, varying the number of tilts from 5 to 121 and the angular range from Δθ = ±10º to ±60º (**Fig. 1c, Extended Data Fig. 1, Supplementary Video 1**). We found that with as little as 21 tilts between Δθ = ±30º, unmyelinated axons, chemical synapses gap junctions and clathrin-coated, spinule-like protrusions between glia and neurons were identifiable (**Fig. 1d, Extended Data Figs. 2, 3a, Supplementary Videos 1-2**). Moreover, reconstructing the same regions with fully-sampled ET could unambiguously validate these structures with additional detail. These results suggest we can perform EM connectomic experiments with limited-tilt ET using ∼50-fold fewer images than fully-sampled ET on sections ∼20-fold thicker than typical EM samples.

**Figure 2.**
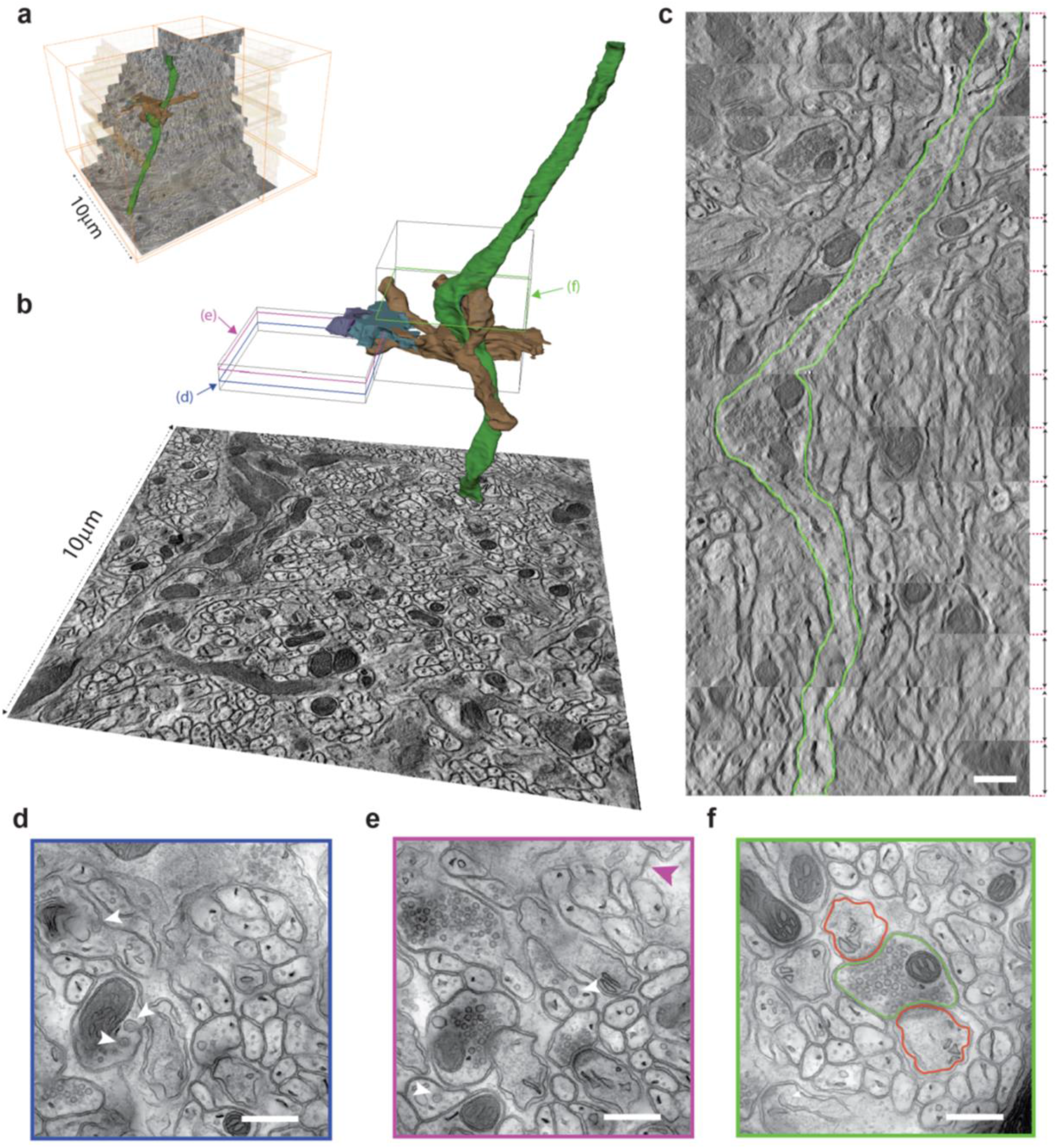
Examples of ET volume stitching, segmentation, and multi-resolution embedding. **(a)** ET images from 15 sequential 600 nm thick sections of mouse cerebellar cortex aligned in Z, showing orthogonal x,y, and z re-slices. Single axis tomography was conducted at low magnification (3.3k). **(b)** Segmentation of a granule cell parallel fiber axon (green) synapsing onto two Purkinje cell spines (brown), astrocytes (purple and blue) and spinules. Boxes outline the areas where multi-axis tomography at higher magnification (11k) was implemented. **(c)** Axon from (a-b, green) segmented (green outline) across 15 aligned 600 nm thick sections projected in an X-Z slice through the volume. Scale bar, 500 nm. **(d)** Multi-axis, high magnification (11k) tomogram featuring spinules (white arrowheads). Scale bar, 500 nm. **(e)** Multi-axis, high magnification (11k) tomogram featuring gap junction (purple arrowhead) and spinules (white arrowheads). Scale bar, 500 nm. **(f)** Multi-axis, high magnification (11k) tomogram featuring the synapse from (a-c). Green outline – parallel fiber axon. Red outlines – Purkinje cell spines. Scale bar, 500 nm.

To quantitatively evaluate image quality across ET acquisition parameters and section thicknesses, we evaluated limited-tilt tomograms against fully-sampled ET reconstructions of the same FOV, and used image (real-space) correlations and Fourier Shell Correlations (FSC)^20^ to compare image quality across different sampling parameters. While the FSC resolution is not a direct measure of resolving power, it offers a resolution estimate that allows quantitative comparisons across imaging parameters. We found that correlations and FSC resolution rapidly improve with relatively few projections, and that FSC resolutions <15 nm can be achieved with parameters compatible with high-throughput imaging (**Fig. 1e**, 11.8 ± 1.3 nm for 500 nm thick, 12.7 ± 0.9 nm for 750 nm thick, 14.8 ± 0.7 nm for 1000 nm thick, mean ± IQR for 21 projections within Δθ = ±30º, **Extended Data Figs. 1-2, 3a-c, 4**). Furthermore, we found that much higher FSC resolutions can be achieved on the same thick sections at higher magnifications (**Fig. 1f**, 4.2 ± 0.1 nm for 500 nm thick, 4.4 ± 0.1 nm for 750 nm thick, 4.6 ± 0.1 nm for 1000 nm thick, mean ± IQR for 21 projections within Δθ = ±30º). We also examined the effect of the angular tilt range, and discovered that for limited-tilts, FSC resolution and correlation-based quality measures are generally highest between Δθ = ±20º to ±30º (**Fig. 1g, Extended Data Figs. 1-2, 3a-c, 4**). This suggests that high-tilt projections may not provide much additional benefit for thick samples in a high-throughput imaging scheme.

We also tested whether semi-thick section ET is feasible at lower accelerating voltage, which may allow for greater accessibility. We found chemical synapses within in a 750 nm thick section were still identifiable at 200 kV with 21 tilts between Δθ = ±30º (**Extended Data Fig. 3d**); however, as expected with lower accelerating voltages, image intensity degrades more quickly at higher angles or thick samples because a larger fraction of electrons are absorbed by the sample.

To obtain volumes large enough for connectomics, multiple tomograms must be stitched together and segmented to map neuronal circuits. Mosaicking, or stitching montages of tomograms within the same physical section has been demonstrated previously^16,21,22^. Here, we aligned and stitched across serial sections, combining fifteen 600 nm thick sections to form a contiguous ∼10 µm × 10 µm × 9µm volume (**Fig. 2, Supplementary Videos 3-4**). To demonstrate that the image quality is sufficient for connectomic analysis, we reconstructed granule cell parallel fiber axons and their synapses onto Purkinje cell dendrites (**Fig. 2b, Supplementary Videos 5-7**). We then collected high resolution (11k) tomograms from subvolumes of the same area, to reveal the same structures in fine detail (**Supplementary Video 7**). These results demonstrate that sectioning at these thicknesses is sufficiently lossless to permit 3D reconstruction of thin axons or synaptic structures, and that limited-tilt ET of semi-thick serial sections is a viable approach for large- and multi-scale connectomics.

## Discussion

High-throughput ET is poised to transform connectomics by providing several key advantages. One inherent strength of the method is its potential to streamline data acquisition. Not only does it reduce the number of serial sections necessary to cut, process, and handle, but thicker sections are also more robust. Another advantage is that it permits high resolution imaging to resolve cellular structures beyond current connectomic datasets. We anticipate that machine learning techniques for limited-tilt tomographic reconstructions will further reduce the number of projections needed per volume. Increasing imaging throughput will allow comparison of connectomes across multiple individuals, enabling experiments between different treatment groups, conditions, or perturbations.

Key challenges remain to scale ET for whole mammalian brain connectomics. Achieving automation in sample positioning across tens of thousands of serial sections while maintaining eucentricity is critical and will require innovation. One way to implement large-scale serial section ET is combining automated tape-based sample handling^10,16,23^. We propose three tape-based ET acquisition strategies to optimize and balance throughput with sample risk (**Supplementary Table 1**). Alternatively, custom large grids (so called Gondola or Super grids) provide a grid-based alternative where volumes as large as a whole adult *Drosophila melanogaster* brain can be collected on a single grid (assuming ∼1 μm sections). Additionally, although higher voltage microscopes are less abundant and more challenging to maintain, we show that ET for connectomics may also be possible on 200 kV microscopes (**Extended Data Fig. 3d**), which are more accessible, but are more susceptible to electron beam absorption and radiation-induced damage. Moreover, although we demonstrated ET with semi-thick (500-1000 nm) sections, integrating additional features such as most probable loss imaging, may enable higher resolution with even thicker samples^24^. Finally, development of advanced electron optics may allow users to rapidly tilt the electron beam, minimizing the overhead required to physically tilt specimens, which would dramatically increase ET throughput.

Multiscale, sample-preserving ET permits extremely high-resolution imaging of volumes of interest *in situ*, which will enable discovery of more intricate and detailed intercellular interactions within the context of larger, lower resolution connectomes. We demonstrate the ability to visualize key structures for intercellular information transfer including electrical synapses and spinule-like structures that appear to be involved in trans-endocytosis between different cell types^12^. We anticipate high-resolution ET imaging will enable discovery of additional cellular structures and mechanisms of information transfer with unprecedented detail, moving us closer to a comprehensive understanding of neuronal circuits.

## Methods

### Experimental animals

Animal procedures were performed in accordance with the National Institutes of Health and Institutional Animal Care and Use Committee (IACUC) guidelines at Harvard Medical School (protocol IS00000124-6) and UCSD (protocol SO6211). Mice (*Mus musculus*) were P36 C57BL/6 or 11 week old 4BALB/cAnNHsd male mice housed on a normal light–dark cycle with an ambient temperature of 18– 23 °C with 40–60% humidity.

### Sample preparation

Mice were perfused transcardially by flushing with Ringer’s solution, followed by perfusion with 0.15M cacodylate buffer (Ted Pella, #18851) containing 2.5% glutaraldehyde (EMS, #18220), 2% freshly made paraformaldehyde (EMS, #19202), and 2mM CaCl_2_ at 37°C. Brains were dissected and cerebellums were separated and placed in vials on ice containing cold fixative for at least two hours. 100 um thick cerebellum tissue sections were cut using a Leica VT 100E vibratome. The buffer is kept at 4°C and ice is packed around the vibratome boat to keep tissue and buffer at 4°C. The sliced tissues were fixed overnight at 4°C. The next day, slices were washed for 5 x 5 minutes in 0.15 M sodium cacodylate buffer containing 2 mM CaCl_2_ at 4°C on ice. The slices were post-fixed with 2% osmium tetroxide (EMS, #19150)-0.8% potassium ferrocyanide solution in 0.15 M sodium cacodylate buffer containing 2 mM CaCl_2_ for 1 hour at 4°C on ice with occasional agitation. Slices were washed with ddH2O 5 x 5 minutes at 4°C on ice and were placed in aqueous 2% uranyl acetate (EMS, #22400) overnight at 4°C. The following day, slices were washed with ddH2O 5 x 5 minutes at 4°C. Slices were dehydrated using ice cold solutions of 50%, 70%, 90%, 100% EtOH, 5 minutes each at 4°C, and at room temperature in solutions of 100% EtOH, 100% EtOH, 1:1 dry acetone – 100% EtOH, and dry acetone for 5 minutes each. Slices were infiltrated in either Epon 812 (EMS, # 14120) or LX-112 resin (Ladd Research, #21311). The infiltrate was 50% resin:acetone overnight, the next day 75% resin:acetone for 2 hours, 90% resin:acetone for 2 hours and 100% resin overnight. Slices were placed in fresh 100% resin 2 x 3 hours. Slices were mounted on microscope slides coated with liquid release agent (EMS, #70880) in 100% resin and cured at 60°C for 2 days. After polymerization of resin, embedded cerebellum samples were cut and glued onto mounting cylinders (Ted Pella, #10580). The mounted slices were trimmed and sectioned using a Leica Ultracut UCT ultramicrotome and a Diatome Ultra 45° diamond knife (Diatome, #40-US). Cerebellum slices embedded in Epon 812 were sectioned at 0.5, 0.75, 1.0, 1.25, 1.50, 1.75 and 2.0 µm thick. Each section was mounted individually onto a Luxel copper slot grid (LUXfilm, # C-S-M-L). Cerebellum slices embedded in LX-112 were serially sectioned at 0.6 µm thick and mounted as a group of four on six Luxel copper slot grids for a total of serial 24 sections. Finally, a mixture of three different size colloidal gold nanoparticles (5 nm, 20 nm and 50 nm) was spread on both surfaces of each sample grid.

### TEM Tomography

Tomography experiments were conducted on a Titan Halo (ThermoFisher Scientific) transmission electron microscope (TEM) operating mainly at 300 kV, and at a lower voltage value (200 kV) in a few instances. Transmitted electrons were collected on a direct detector DE64 composed by an array of 8k × 8k pixels (Direct Electron, LP). Throughout this work, the microscope magnification ranged from 3.3k to 11k, the original micrograph pixel size varying from 1.23nm to 0.36nm; For practical reasons, we only used a four times binned version of the data, with pixel size values spanning then from 4.91nm to 1.44nm.

We predominantly adopted a single tilt acquisition scheme where the sample is rotated in the electron microscope beam at fixed angle increment around a unique sample axis. The angle increment was chosen to be very fine (0.25°) while rotations were between -60 and +60 degrees, allowing us to easily extract any subset of tilts to subsample the total set of projections. In a few cases, we also introduced a more sophisticated protocol consisting of acquiring multiple tilt series (typically 4) shot on the same sample area but at evenly spaced angles and leading to better 3D reconstructions^18^. It provided us (i) with a benchmark for the most optimal reconstruction to achieve for a given magnification and sample thickness, and (ii) allowed us to assess quantitatively what resolution to expect in a reconstruction by comparing two volumes arising from different tilt series using a Fourier shell correlation.

We used identical parameters for angle span and increment in the full data acquisition scheme, regardless of the sample thickness under consideration. Data was acquired using SerialEM (https://bio3d.colorado.edu/SerialEM/).

### Tomographic Reconstruction

To assess which biological features can be discerned when using a specific acquisition scheme for tomography, we generated an extensive set of 3D reconstructions for all the regions of interest, each reconstruction built from a distinct specimen orientation pattern. This allowed us to systematically characterize a large portion of the reconstruction space and pinpoint the most efficient scheme for connectomics.

To parameterize acquisition space, we started with a constant increment δθ of the full angle range Δθ (=2 θ_max_), and the number of projections N in a tilt series were selected from {±10, ±20, ±30, ±40, ±50, ±60} and {5, 11, 17, 21, 33, 121}, corresponding to a value δθ =Δθ (N-1). Tilt angles (Δθ, N), that is the set - θ_max_+ *i* δθ with 0 ≤ i ≤ N, were mapped to their closest values in the full acquisition (Δθ_0_=±60, N_0_=481), to specify which projections to include in the reconstruction.

The micrographs were aligned for the entire data set (Δθ_0_=±60, N_0_=481) with a single bundle adjustment procedure, and the result shared between the different data acquisition schemes (Δθ, N). Likewise a global alignment procedure was used if the reconstruction involved multiple tilts (see FSC calculations).

Tomograms were generated using filtered back-projection unless otherwise specified. In some cases, we used a weighted iterative reconstruction approach that provided higher quality volumes. This was used in **Figure 2**.

### Image Analysis

#### Fourier Shell Correlation

Fourier Shell Correlation (FSC)^20^ was performed to estimate the resolution of tomographic reconstructions. FSC involves calculating correlations between two independent image volumes. For this, we obtained two tilt series with the rotation axis oriented 90-degrees relative to each other (dual-tilt axis tomography). We will refer to these two tilt series as “a” and “b” series. We aligned the two tilt series together, but reconstructed two separate volumes from each single-tilt series. For one series, we reconstructed a high-quality reference volume using the full series (481 projections over Δθ = ±60°). For the other series, we reconstructed test volumes using a limited number of projections and angular range. FSC was performed by measuring the normalized cross-correlation coefficient between the reference and test volumes over corresponding shells in Fourier space. The resolution was determined by the intersection between the measured FSC and the 1-bit threshold as a function of spatial frequency^20^. The more conservative 1-bit threshold was used (as opposed to the more conventional ½-bit threshold) because we are interpreting the FSC resolution to be the resolution of the limited tilt test volume, rather than the combination of the test and reference volumes^25^. We computed FSC in a symmetrical manner: with the “a” series as reference and “b” as test, and vice versa, then averaged the for the final result. For the FSC analysis presented here, it was not acceptable to simply divide a single-tilt series into even and odd projections because the missing wedge artifacts, which are similar in both reconstructions, create spurious correlations. When dual tilt series are compared, the missing wedge artifacts are not equivalent, reducing the likelihood of spurious correlations.

FSC measurements were performed using custom code based on functions provided in the Toupy package^26^. To account for variability of image quality, the reconstructed volumes were divided into chunks of 45^3^ voxels, and FSC was performed on each chunk independently. Thus, the distribution of FSC resolution values across chunks captures the variability within the reconstructed volumes. The size of the chunks was chosen such that an integer number of chunks comprises most of the thickness (>80%) of the reconstructed sections (e.g. 500 nm, 750 nm, and 1000 nm). The outer 79 voxels in the remaining two axes were excluded to avoid artifacts related to alignment of the tilt series, and to ensure that the volumes can be divided into an integer number of chunks.

It is important to note that these FSC resolutions are an estimate of approximate resolution, rather than a direct measurement of resolving power. The resolution in limited-tilt reconstructions is anisotropic, where the in-plane resolution is usually superior to the axial resolution, but 3D FSC measurements convolve resolution along different dimensions together. Moreover, parameters in the FSC measurements – most notably the threshold – can have a direct effect on the FSC resolution values^20^, so care must be taken in comparing these results to other studies. Nevertheless, while the absolute values of FSC resolution should be interpreted cautiously, the relative values between equivalent FSC measurements allows quantitative comparison across different levels of subsampling.

#### Image Correlations

Real-space image correlations were calculated from dual-tilt series similarly to FSC, except Pearson correlations were calculated in real-space instead of in Fourier-space. From one tilt series, we reconstructed a high-quality reference volume using the full series (481 projections over Δθ = ±60°). For the other series, we reconstructed test volumes using a limited number of projections and angular range. The two independent volumes were divided into 45^3^ voxel chunks, and the Pearson linear correlation between the test and reference volumes was calculated for corresponding chunks. Correlations were computed in a symmetrical manner: with the “a” series as reference and “b” as test, and vice versa, then averaged for the final result. Thus, the correlation values are an approximation of the similarity between the limited-tilt reconstruction and the “ground truth” volume. It is likely that these correlations are under-estimates, because the reference volumes contain noise and missing wedge artifacts. Nevertheless, the goal of these measurements is to make a relative comparison of the correlations as the number of projections and angular range are varied. FSC and image correlation measurements for tomograms acquired at 3.3k magnification included 2-3 different tomograms sampled from different parts of the sample for each thickness (7 in total), whereas 1 tomogram per thickness was used for 11k magnification (3 in total).

### Serial Tomogram Alignment

To stack and align the tomograms, we first estimated their normal direction (Z direction). This can be accurately derived from the gold markers positions if those particles are well distributed on the specimen surfaces. Tomogram boundaries are then identified by the maximum values in the gradient norm along Z. Next, we used 2D affine maps to register the surfaces between two adjacent tomograms. Their rotation was first evaluated by registering the images of adjacent surfaces in polar coordinates. The remaining residual part of the transform was then obtained by correlating patches from one surface image to its rotated counterpart. To align the entire serial stack, the relative maps between consecutive tomograms were composed and the final output transform was applied to each tomogram. Any slow-moving drift or shape modification along the Z direction was canceled. The mouse cerebellum serial stack was aligned from 15 serial sections collected on three different grids.

There was no need to digitally flatten the reconstructions during this stack and align procedure. Acquiring tomographic data at low magnification (3.3k) does not require a large beam surface density on the sample to produce micrographs with a satisfactory contrast. This also helps prevent out of plane distortions, which are difficult to correct.

### Segmentation & Image Rendering

Contrast of different 3D reconstruction figure snapshots were adjusted for clarity. Low spatial frequencies in the DE64 tomograms were adjusted to be consistent with volumes obtained from a CCD detector. This consisted of calculating the average of a tomogram and its Gaussian blurred counterpart (with 3D kernel value around 1 pixel) and re-scaling the image intensity histogram so that 98% of voxels fill the dynamic range.

We manually segmented a few biological features within the large serial stack of the mouse cerebellum (**Fig. 2**), to illustrate the feasibility of tracking them across consecutive sections. Those include a neuronal axon, dendrite, synapse, astrocytes, vesicles, as well as other organelles. The segmentation was built upon the fully-sampled version of the stack (acquired at 3.3k magnification and a 4.91nm pixel size) so the smallest components (synaptic vesicles) could still be delineated. Segmentation was done using 3dmod (https://bio3d.colorado.edu/imod/doc/3dmodguide.html).

Notably, the segmented unmyelinated axon remains continuous throughout the entire volume (**Fig. 2B**). In this view, we extracted the surrounding of the neuron and projected it on the stack front view (which defines here the x and y axes). To be more precise, we extracted the voxel values located on the surface that tracks the neuron changes along the y direction but remains constant in the x direction. This procedure was effective because the neuron path is mostly monotonic in the y direction, with no complicated folding. To compute the surface of interest, the segmented neuron was first reduced into a single filament by directly skeletonizing its segmentation.

The views in **Figure 2B** and **Supplementary Video 4** make it clear that any loss of material in between sections appears negligible and would not preclude connectomics studies. A more refined tomography process was also implemented on the previously segmented synapse at a higher magnification (11k / 1.44nm pixel size) and with a full 4-tilt approach; it confirmed that split vesicles can be tracked across sections, and any loss of material can be estimated to <10 nm.

Amira (ThermoFisher Scientific) was used for rendering and animations. For renderings and animations including tomograms with different resolutions, we used an approach like the serial tomogram stacking procedure described above to superpose the different volumes.

## Acknowledgements

This work was supported by the NIH (RF1MH129261 to M.E. and W.C.A.L. and K99EB032217 to A.T.K.).

## Author Contributions

Conceptualization: A.T.K., S.P., S.T.P., M.E. W.C.A.L; Sample preparation: M.M., M.K., Data acquisition, processing, and visualization: S.P; Data analysis: A.T.K., S.P.; Cell segmentation: K-Y.K.; Manuscript writing and editing: A.T.K., S.P., W.C.A.L. with input from other authors.

## Extended Data Figures

**Extended Data Fig. 1.**
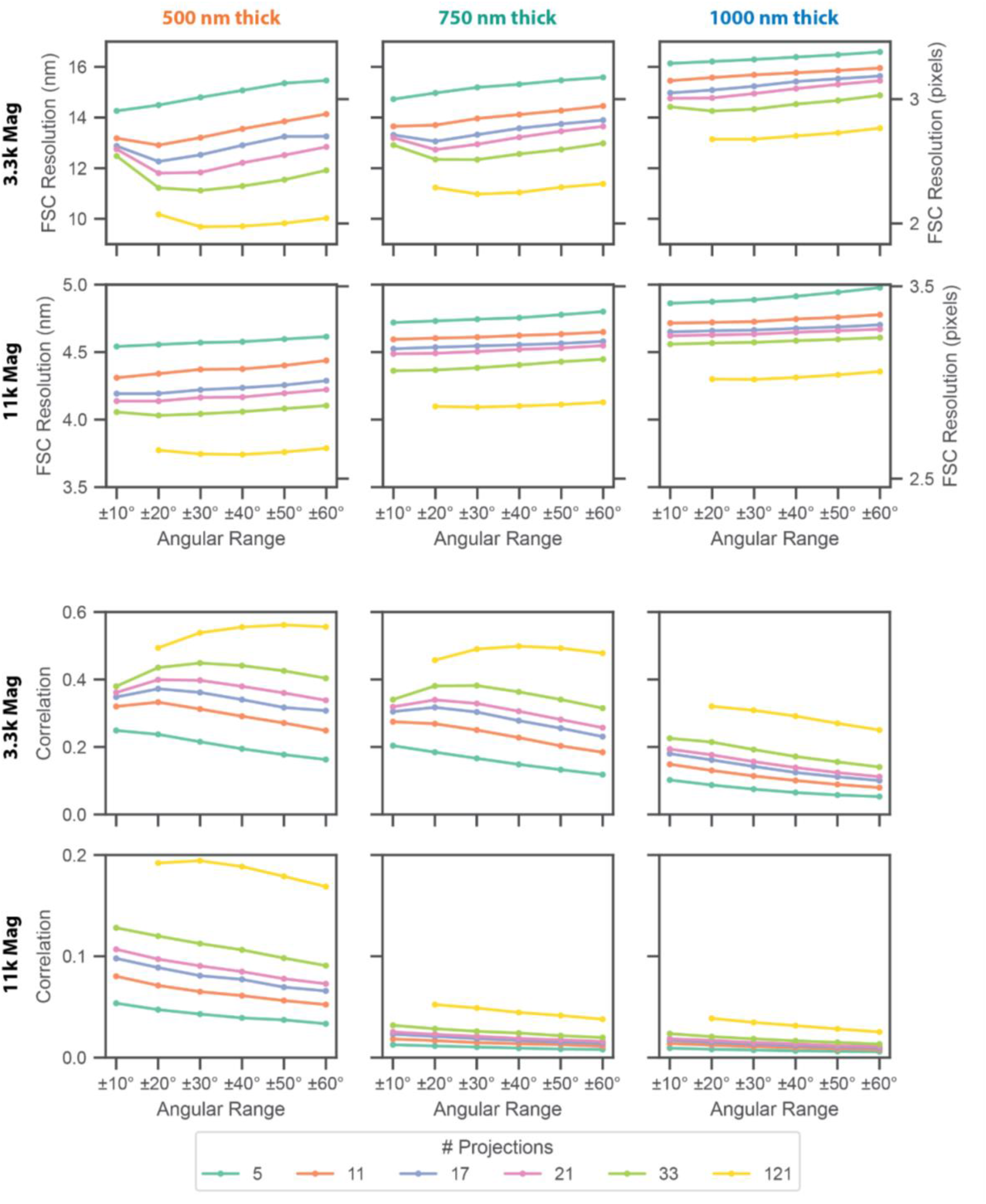
Quantifications of ET reconstruction quality at 300 kV. Quantifications of mean FSC resolution (top 2 rows) and correlations with reference images (bottom 2 rows) for limited-tilt ET reconstructions (Methods). Plots are arranged by sample thickness (columns: left: 500 nm, middle: 750 nm, right: 1000 nm) and magnification (1st and 3rd row: 3.3k, 2nd and 4th row: 11k). The number of projections is indicated by line and marker color (legend at bottom).

**Extended Data Figure 2.**
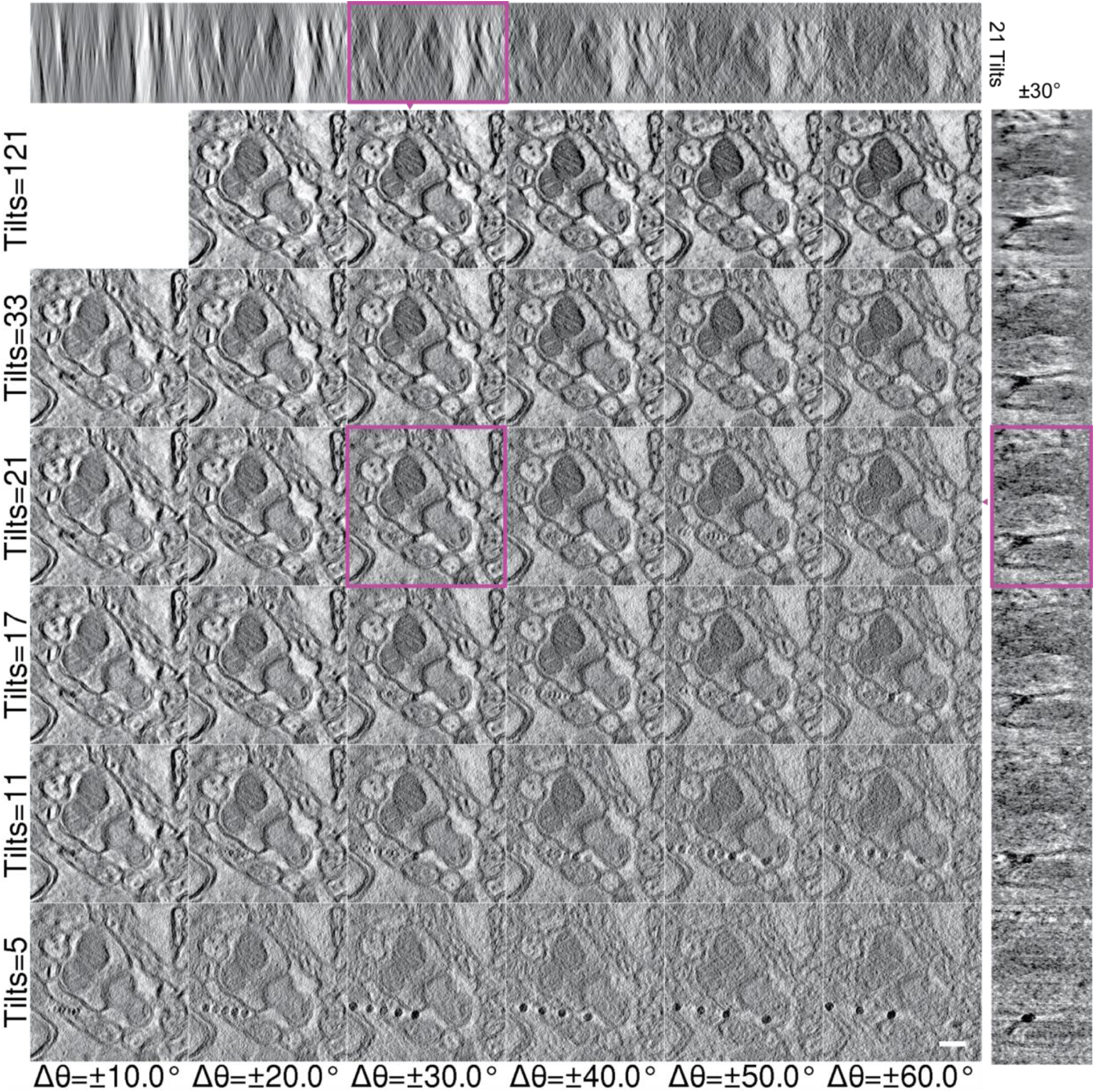
ET operating point image matrix for connectomics. A mosaic of ET images across different tilt ranges (x-axis) and number of tilt angles (y-axis, evenly sampled over the tilt range) of an excitatory synapse in the cerebellar molecular layer. *(top)* X-Z and *(right)* Y-Z virtual slices through the tomographic volume at different operating points. The parameter set (Δθ = ±30, N = 21) outlined in purple is relevant to connectomic. Scale bar, 250 nm.

**Extended Data Fig 3.**
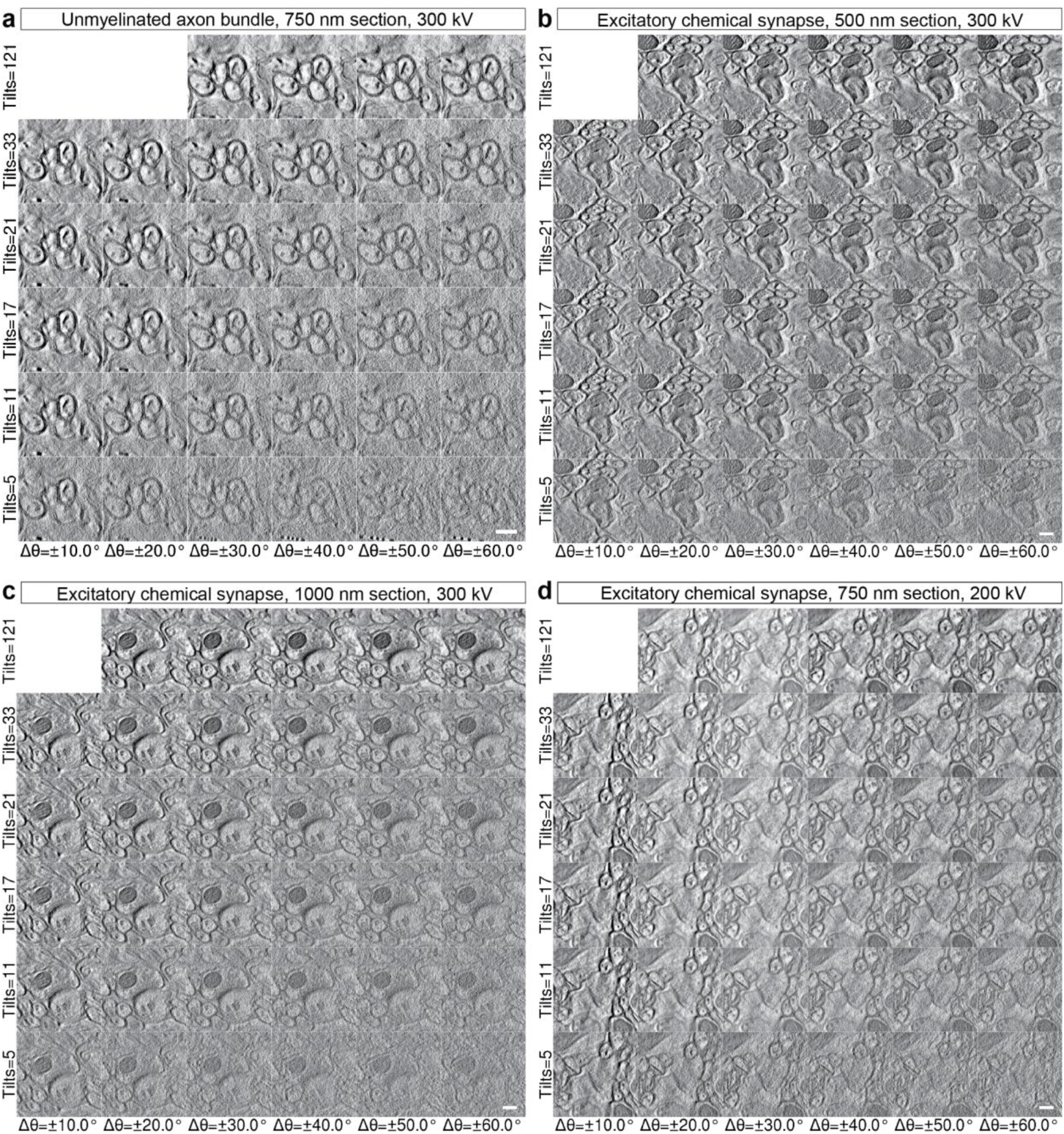
ET operating point parameterization at lower voltage and differing section thicknesses. **(a)** Parameterization mosaic of ET images across different tilt ranges (x-axis) and number of tilt angles (y-axis, evenly sampled over the tilt range) of an unmyelinated axon bundle. Scale bars, 250 nm **(b)** Similar to Extended Data Fig. 2a with 500 nm thick sections. **(c)** Similar to Extended Data Fig. 2a with 1000 nm thick sections. **(d)** Parameterization mosaic of ET images of an excitatory synapse (similar to Extended Data Fig. 2a) at 200 kV.

**Extended Data Fig. 4.**
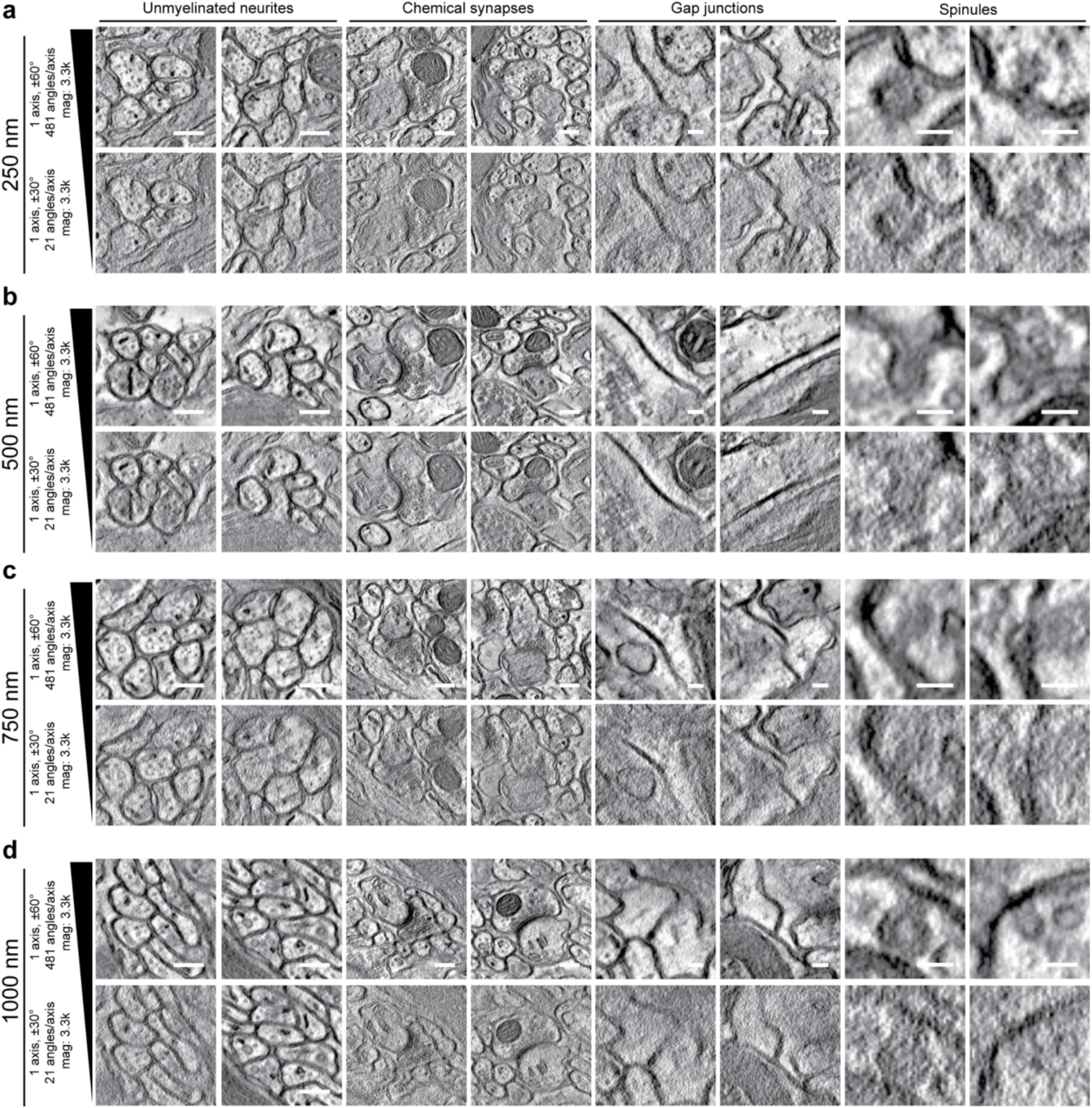
Biological features in ET with differing section thickness. **(a)** Similar to Fig. 1d with 250 nm thick sections, full (top row) and limited (bottom row) single-axis tomography. Scale bars: Unmyelinated neurites and Chemical synapses, 250 nm; Gap junctions, 100 nm; Spinules, 75 nm. **(b)** Similar to (a) from 500 nm thick sections. **(c)** Similar to (a) from 750 nm thick sections. **(d)** Similar to (a) from 1000 nm thick sections.

## Supplementary Information

**Supplementary Video 1. Example ET image volumes at different tilt ranges and sampling**.

Fly-through of mosaics for ET volumes across different tilt ranges (x-axis) and number of tilt angles (y-axis, evenly sampled over the tilt range). Starting with a virtual 8k × 8k pixel section from the molecular layer of an adult mouse cerebellar cortex, zooming in to an operating point (121 tilts, angle range Δθ = ±60°), moving to a different operating point (21 tilts, angle range Δθ = ±30°), followed by flying through the same unmyelinated neurite bundle with each operating point mosaiced. Note, differences in data quality with different tilt ranges and angles.

**Supplementary Video 2. Example ET of axons at different tilt ranges and sampling**.

Fly-through mosaic of ET images across different tilt ranges (x-axis) and number of tilt angles (y-axis, evenly sampled over the tilt range) of an unmyelinated bundle of axons in the cerebellar molecular layer.

**Supplementary Video 3. Example high-resolution ET image volume**.

Fly-through of an aligned region through 15 sequential 600 nm thick section tomograms from the molecular layer of the adult mouse cerebellar cortex. Scale bar, 500 nm.

**Supplementary Video 4. Example aligned ET volume**.

Fly-through of an aligned region through 15 sequential 600 nm thick section tomograms re-sliced in Z.

**Supplementary Video 5. Neuron segmentation from an ET volume**.

Segmentation of a granule cell parallel fiber axon (green) synapsing onto two Purkinje cell spines (off white) with their associated mitochondria (yellow), spine apparati (cyan), and synaptic vesicles (small red spheres) emerging from aligned tomograms of 15 sequential 600 nm thick section tomograms.

**Supplementary Video 6. Following axons across aligned semi-thick section ET**.

Fly-through following the granule cell parallel fiber axon (green outline) segmented in **Figure 2** and **Supplementary Video 5** through 15 aligned tomograms with (*left*) 21 tilts between Δθ = ±30º and (*right*) 241 tilts between Δθ = ±60º. Scale bar, 500 nm.

**Supplementary Video 7. Example alignment of multiple-axis and limited-tilt ET volumes**.

Video of a high-resolution, multi-axis tilt ET aligned into a larger, lower-resolution ET volume with neurons segmented as in **Figure 2**.

**Supplementary Table 1.**
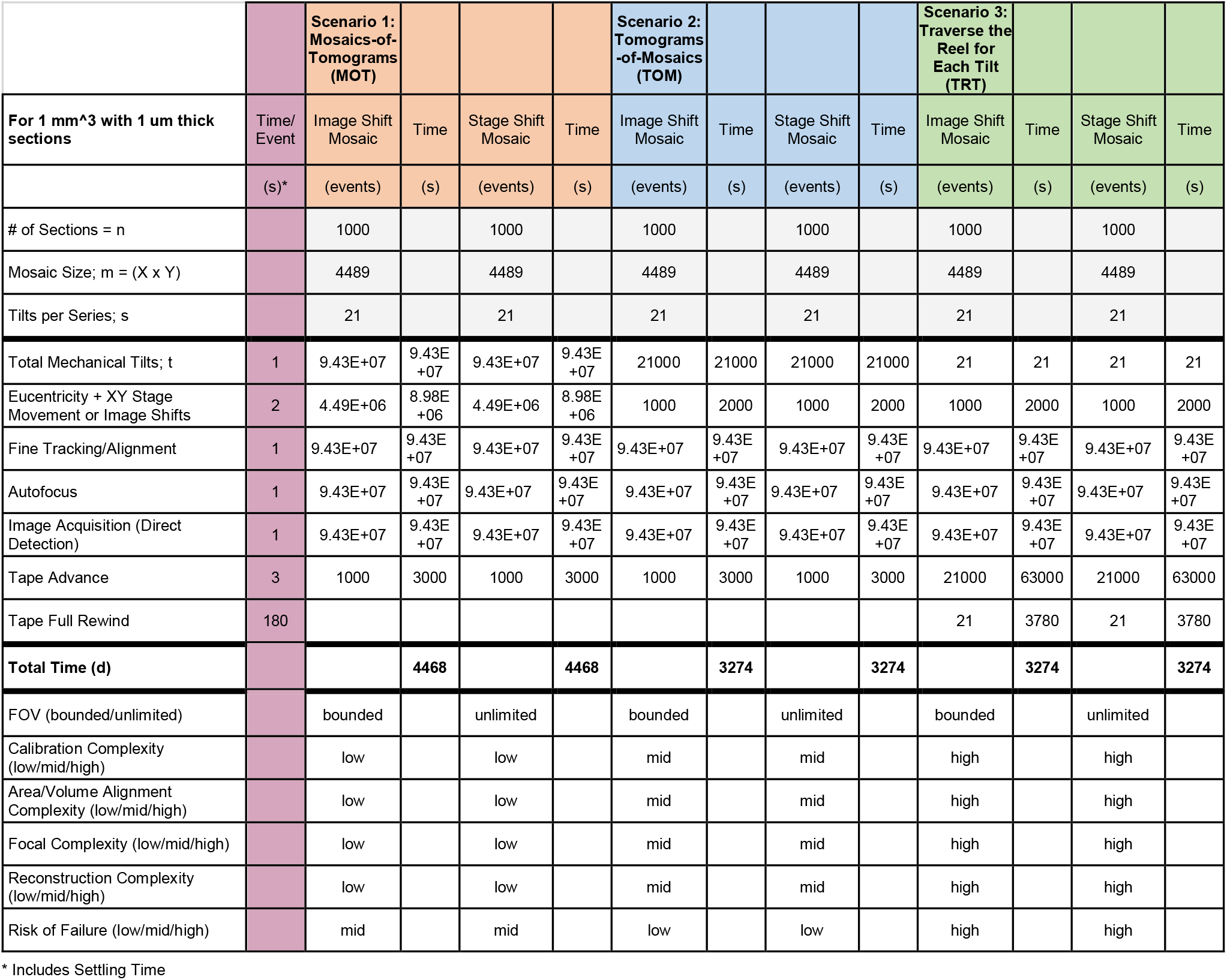
Acquisition strategies for scaling ET. Tape-based ET acquisition strategies to optimize and balance throughput with sample risk: (1) Mosaics-of-Tomograms (MOT): Capture angular projections for each subtomogram at specific x,y positions across each section’s region of interest (ROI), utilizing either image shift or physical stage shifting. Despite its apparent simplicity in implementation and alignment, this method involves numerous mechanical tilts, increasing the overhead of movement and sample settling. (2) Tomograms-of-Mosaics (TOM): For each tilt angle, acquire an ROI-spanning montage, before tilting and mosaicing again. This reduces mechanical tilts; however, it requires intricate stage calibration to keep the sample in focus while moving in x & y (eucentricity maintenance) potentially impacting overall efficiency. (3) Traverse the Reel for each Tilt (TRT): Acquire section montages for an angle across every section on the reel, then change angle, and image the whole reel in reverse before changing angles again. This approach minimizes mechanical movement, but increases the risk of sample damage due to frequent tape traversals. This method demands significant calibration efforts for imaging, reconstruction, and alignment. Overall, these strategies provide a framework for future advancements in large-scale ET.

## References

1. Eichler, K. et al. The complete connectome of a learning and memory centre in an insect brain. Nature 548, 175–182 (2017).

2. Nguyen, T. M. et al. Structured cerebellar connectivity supports resilient pattern separation. Nature 613, 543–549 (2023).

3. Ohyama, T. et al. A multilevel multimodal circuit enhances action selection in Drosophila. Nature 520, 633–639 (2015).

4. Kuan, A. T. et al. Synaptic wiring motifs in posterior parietal cortex support decision-making. Nature (2024) doi:10.1038/s41586-024-07088-7.

5. Azevedo, A. et al. Tools for comprehensive reconstruction and analysis of Drosophila motor circuits. bioRxiv 2022.12.15.520299 (2022).

6. Lesser, E. et al. Synaptic architecture of leg and wing motor control networks in Drosophila. bioRxiv (2023) doi:10.1101/2023.05.30.542725.

7. Eberle, A. L. et al. High-resolution, high-throughput imaging with a multibeam scanning electron microscope. J. Microsc. 259, 114–120 (2015).

8. Xu, C. S. et al. Enhanced FIB-SEM systems for large-volume 3D imaging. Elife 6, (2017).

9. Hayworth, K. J. et al. Gas cluster ion beam SEM for imaging of large tissue samples with 10 nm isotropic resolution. Nat. Methods 17, 68–71 (2020).

10. Phelps, J. S. et al. Reconstruction of motor control circuits in adult Drosophila using automated transmission electron microscopy. Cell (2021) doi:10.1016/j.cell.2020.12.013.

11. Fields, R. D. & Stevens-Graham, B. New insights into neuron-glia communication. Science 298, 556–562 (2002).

12. Spacek, J. & Harris, K. M. Trans-endocytosis via spinules in adult rat hippocampus. J. Neurosci. 24, 4233–4241 (2004).

13. Soto, G. E. et al. Serial Section Electron Tomography: A Method for Three-Dimensional Reconstruction of Large Structures. Neuroimage 1, 230–243 (1994).

14. Phan, S. & Lawrence, A. Tomography of large format electron microscope tilt series: Image alignment and volume reconstruction. in 2008 Congress on Image and Signal Processing vol. 2 176–182 (IEEE, 2008).

15. Berlanga, M. L. et al. Three-dimensional reconstruction of serial mouse brain sections: solution for flattening high-resolution large-scale mosaics. Front. Neuroanat. 5, 17 (2011).

16. Phan, S. et al. TxBR montage reconstruction for large field electron tomography. J. Struct. Biol. 180, 154–164 (2012).

17. Wan, X. et al. Iterative methods in large field electron microscope tomography. SIAM J. Sci. Comput. 35, S402–S419 (2013).

18. Phan, S. et al. 3D reconstruction of biological structures: automated procedures for alignment and reconstruction of multiple tilt series in electron tomography. Adv Struct Chem Imaging 2, 8 (2017).

19. Wesseling, J. F. et al. Sparse force-bearing bridges between neighboring synaptic vesicles. Brain Struct. Funct. 224, 3263–3276 (2019).

20. Harauz, G. & Heel, M. Exact filters for general geometry three dimensional reconstruction. Optik 73, 146–156 (1986).

21. Mastronarde, D. N., van der Heide, P. A., Morgan, G. P. & Marsh, B. J. Supermontaging: Reconstructing Large Cellular Volumes by Stitching Together Laterally Adjacent Tomograms. Microsc. Microanal. 14, 106–107 (2008).

22. O’Toole, E., van der Heide, P., Richard McIntosh, J. & Mastronarde, D. Large-Scale Electron Tomography of Cells Using SerialEM and IMOD. in Cellular Imaging: Electron Tomography and Related Techniques (ed. Hanssen, E.) 95–116 (Springer International Publishing, Cham, 2018).

23. Consortium, M. et al. Functional connectomics spanning multiple areas of mouse visual cortex. bioRxiv (2021) doi:10.1101/2021.07.28.454025.

24. Bouwer, J. C. et al. The application of energy-filtered electron microscopy to tomography of thick, selectively stained biological samples. Methods Cell Biol. 79, 643–660 (2007).

25. van Heel, M. & Schatz, M. Fourier shell correlation threshold criteria. J. Struct. Biol. 151, 250–262 (2005).

26. Julio Cesar da Silva. Jcesardasilva/toupy: Version 0.2.1. doi:10.5281/zenodo.5139389.

